# Evaluating the cost of pharmaceutical purification for a long-duration space exploration medical foundry

**DOI:** 10.1101/2021.06.21.449300

**Authors:** Matthew J. McNulty, Aaron J. Berliner, Patrick G. Negulescu, Liber McKee, Olivia Hart, Kevin Yates, Adam P. Arkin, Somen Nandi, Karen A. McDonald

**Author notes:** https://cubes.space/.

## Abstract

There are medical treatment vulnerabilities in longer-duration space missions present in the current International Space Station crew health care system with risks, arising from spaceflight-accelerated pharmaceutical degradation and resupply lag times. Bioregenerative life support systems may be a way to close this risk gap by leveraging *in situ* resource utilization (ISRU) to perform pharmaceutical synthesis and purification. Recent literature has begun to consider biological ISRU using microbes and plants as the basis for pharmaceutical life support technologies. However, there has not yet been a rigorous analysis of the processing and quality systems required to implement biologically-produced pharmaceuticals for human medical treatment. In this work, we use the equivalent system mass (ESM) metric to evaluate pharmaceutical purification processing strategies for longer-duration space exploration missions. Monoclonal antibodies, representing a diverse therapeutic platform capable of treating multiple space-relevant disease states, were selected as the target products for this analysis. We investigate the ESM resource costs (mass, volume, power, cooling, and crew time) of an affinity-based capture step for monoclonal antibody purification as a test case within a manned Mars mission architecture. We compare six technologies (three biotic capture methods and three abiotic capture methods), optimize scheduling to minimize ESM for each technology, and perform scenario analysis to consider a range of input stream compositions and pharmaceutical demand. We also compare the base case ESM to scenarios of alternative mission configuration, equipment models, and technology reusability. Throughout the analyses, we identify key areas for development of pharmaceutical life support technology and improvement of the ESM framework for assessment of bioregenerative life support technologies.

## 1. Introduction

### 1.1 The need for a pharmaceutical foundry in space

Surveying missions to Mars, like the InSight lander^1^ launched in 2018 and Perseverance rover^2^ in 2020, directly support the objectives of NASA’s long-term Mars Exploration Program^3^: an effort to explore the potential for life on Mars and prepare for human exploration of Mars. The maturation of the program requires redefining the risks to human health as mission architectures transition from the current Earth Reliant paradigm used on the International Space Station (ISS) to the cislunar space Proving Grounds and finally to deep-space Earth Independent mission architectures, as defined in NASA’s Journey to Mars: Pioneering Next Steps in Space Exploration^4^.

Human missions to Mars will be Earth Independent, meaning there will be very limited emergency evacuation and re-supply capabilities along with substantially delayed communications with the Earth-based mission team. The NASA Human Research Roadmap (https://humanresearchroadmap.nasa.gov/Risks/) currently rates most human health risks, which include ‘risk of adverse health outcomes & decrements in performance due to inflight medical conditions’ and ‘risk of ineffective or toxic medications during long-duration exploration spaceflight’, as either medium or high risk for a Mars planetary visit/habitat mission. Risk ratings are based on failure mode and effects analysis and on hazard analysis using dimensions of severity, occurrence, and detectability. A recent review highlights the current understanding of the primary hazards and health risks posed by deep space exploration as well as the six types of countermeasures: protective shielding, biological and environmental temporal monitoring, specialized workout equipment, cognition and psychological evaluations, autonomous health support, and personalized medicine^5^.

Of these countermeasures, it could be argued that medicine is the most crucial and least advanced towards mitigating space health hazards. There is very limited information on pharmaceutical usage and therapeutic efficacy (i.e., pharmacokinetics, pharmacodynamics) in spaceflight or in a Mars surface environment^6^. Furthermore, flown stores of pharmaceuticals face two additional barriers: 1) radiation-accelerated degradation^7^, and 2) addressing a myriad of low occurrence and high impact health hazards without the ability to fly and maintain potency of therapeutics for all of them. In these circumstances, it is often more beneficial to build robustness to these low occurrence health hazards rather than to try to predict them. It is therefore imperative that on-planet and/or in-flight pharmaceutical production be developed to bridge this risk gap.

Pharmaceuticals are produced either chemically or biologically. A recent review of pharmaceutical production for human life support in space compares these two methods, highlighting the need for biological production in order to address many low occurrence and high impact health hazards and further comparing different biological production systems^8^. One major advantage of biological production is the efficiency in transporting and synthesizing genetic information as the set of instructions, or sometimes the product itself, to meet the therapeutic needs for a variety of disease states. The emerging field of Space Systems Bioengineering^9^ encapsulates this need for biological production, of which pharmaceuticals is identified as an important subset.

### 1.2 The bottleneck of space foundries: purification

Biopharmaceuticals must be purified after accumulation with the biological host organism, or cell-free transcription-translation reaction, in order to meet requirements for drug delivery and therapeutic effect^10^. The majority of commercial biopharmaceutical products are administered via intravenous and subcutaneous injection^11^. Biopharmaceutical formulations for injection requires high purity (>95%) product, as impurities introduced directly into the bloodstream can trigger significant immune responses and reduce efficacy^12^.

Downstream processing of biopharmaceuticals is therefore usually a resource-intensive section of overall processing, being cited as high as 80% of production costs for monoclonal antibody (mAb) therapeutics produced using mammalian cell cultures^13^. In addition to the processing burden for biopharmaceutical injectables, there are also often substantial storage costs involving complex supply chain and storage management with stability requirements for factors including temperature, time, humidity, light, and vibration^14^. There are several approaches being pursued to overcome the challenges and costs associated with downstream processing and formulation.

First are the tremendous efforts in process intensification^15^. While the highly sensitive nature of biopharmaceuticals to minor process changes has introduced barriers and complexities to innovation through process intensification that have not been realized in non-healthcare biotechnological industries, there have been significant strides made in the past decade in the areas of process integration^16^, automation^17^, and miniaturization^18,19^.

Another route that researchers are pursuing to reduce downstream processing costs and resources is a biological solution to processing technology. In the same vein that the biopharmaceutical industry sprung out of researchers leveraging the power of biology to produce therapeutically relevant molecules that were inaccessible or excessively costly by means of chemical synthesis, researchers are now also trying to apply that same principle to purifying therapeutically relevant molecules. The simplicity of production, reagents that can be produced using self-replicating organisms, and potential recyclability of spent consumables are significant advantages of biological purification technology for space or other limited resource applications. Examples of primary biological technologies include fusion tags^20^, stimuli-responsive biopolymers^21^, hydrophobic nanoparticles^22^, and plant virus nanoparticles^23,24^.

Lastly, there are vast efforts to establish alternative drug delivery modalities^25^. Other modalities that do not require injection and which might be more compatible to administration in limited resource environments, such as oral consumption, nasal spray, inhalation, and topical application, have long presented challenges in biopharmaceutical stability (e.g., denaturation in stomach acid) and delivery to the active site (e.g., passing the gut-blood barrier) that minimize product efficacy and necessitate costly advanced formulations and chemistries^26^.

A particularly promising drug delivery technique to circumvent downstream processing burdens is to sequester the active pharmaceutical ingredient in the host cells of the upstream production system as a protective encapsulation in order to facilitate bioavailability through oral delivery^27^. It represents an opportunity to greatly lower the cost of *in situ* production of human medicine for a space mission. This technique presumes that the host system is safe for human consumption, and so naturally lends itself to utility in systems such as yeast and plant production hosts. Oral delivery via host cell encapsulation has been recently established as commercial drug delivery modality with the US Food and Drug Administration approval of Palforzia as an oral peanut-protein immunotherapy^28^. However, this solution is not necessarily amenable to the diversity of pharmaceutical countermeasures that may be required, especially for unanticipated needs in which the product may not have been evaluated for oral bioavailability.

### 1.3 Space economics

In 2011, the space shuttle program was retired due to increasing costs, demonstrating that reduction of economic cost is critical for sustaining any campaign of human exploration^29^. Although recent efforts in reducing the launch cost to low earth orbit by commercial space companies have aided in the redefinition of the space economy^30^, the barrier to longer term missions, such as a journey to Mars, is still limited by the extreme financial cost in transporting resources. Additionally, it has been shown that as the mission duration and complexity increases – as expected for a human mission to Mars – the quantity of supplies required to maintain crew health also increases^31^. In the case of meeting the demand for medication, biopharmaceutical synthesis has been proposed as an alternative to packaging a growing number of different medications^8,32^. Assuming that both technologies can meet mission demand, selection of the production-based biotechnology platform will be dependent on its cost impact. It is therefore critical that the cost model of biopharmaceutical synthesis accounts for and minimizes the cost of any and all subprocesses, including those for purification.

The current terrestrial biopharmaceutical synthesis cost model does not align with the needs for space exploration environments. For example, the literature highlights the high cost of Protein A affinity chromatography resin ($8,000 – $15,000/L) and the need to reduce the price^33^. However, the purchase cost of chromatography resin is not nearly as critical in space environment applications where the major costs are more closely tied to the physical properties of the object (mass, volume, refrigeration requirements, etc.), as a result of fuel and payload limitations and the crew time required for operation^34^. The distinct cost models of space and terrestrial biopharmaceutical production may increase the burden of identifying space-relevant processing technologies and may also limit direct transferability of terrestrial technologies without attention given to these areas.

On the other hand, changing incentives structures relating to sustainability and the advent of new platform technologies are rapidly increasing alignment and the potential for technology crossover. For example, companies like On Demand Pharmaceuticals (ondemandpharma.com), EQRx (eqrx.com), and the kenUP Foundation (kenup.eu), initiatives leading to industry adoption of environmental footprint metrics such as E-factor^35^ and Process Mass Intensity (PMI)^36^, and diffusion from the adjacencies of green and white biotechnology^37^ all promote development of accessible and sustainable technologies. As these trends pertain to space-relevant processes, these examples can also be viewed as driving more closed loop systems composed of simpler components.

#### 1.3.1 Reference mission architecture

The evaluation of biopharmaceutical system cost for space applications requires the establishment of a reference mission architecture (RMA) as a means for describing the envelope of the mission scenario and distilling initial technology specifications which relate to the proposed subsystem in question^38^. This RMA can be used to orient and define the specific mission elements that meet the mission requirements and factor into the calculations of cost for deploying biopharmaceutical technologies. Ultimately, the RMA provides the means to determine and compare cost given specification of mission scenarios that utilize the technology in question. We envision developing and integrating biotechnological capabilities back-ended by purification and quality systems into standard methods composed of a series of unit procedures that maintain astronaut health via the Environmental Control and Life Support Systems^39^. In this study, we begin to build towards this vision by proposing a high-level RMA that specifies a biopharmaceutical demand partially fulfilled through biomanufacturing over the course of a defined production window.

#### 1.3.2 Equivalent System Mass (ESM)

In planning for future human exploration missions, technology choices and life-support systems specifications are often evaluated through the metric of the equivalent system mass (ESM)^40^. Driven by the economic factor of cost in dollars required to transport mass into orbit, the ESM framework accounts for non-mass factors such as power, volume, and crew-time by relating them to mass through predetermined equivalency factors. ESM has been used to evaluate the mass of all of the resources of a larger system including water, shielding materials, agriculture and recycle loop closure. Currently, ESM remains the standard metric for evaluating advanced life support technology platforms^41^. In the Space Systems Bioengineering context of realizing a biomanufactory on the surface of Mars^9^, recent advances in extending this metric have been proposed in the form of extended equivalent system mass (xESM) which attempts to address complexities stemming from multiple transit and operations stages, as would be required to support a crewed mission to Mars^42^. The xESM also provides an accountancy for the uncertainties inherent in mission planning. Such advances in the xESM framework aid in the assessment of biopharmaceutical technologies as elements in the context of proposed life support systems given the inherent stochastic nature of human health, especially in a space environment^43^. Here, we leverage the xESM calculations to account for multiple locations across which biopharmaceutical purification is deployed.

## 2. Materials & Methods

### 2.1 Unit procedure selection

#### 2.1.1 Protein A-based affinity capture step

The medical significance of mAbs and the highly developed and specialized purification technology provide a fertile ground for techno-economic feasibility analysis of an ISRU-based pharmaceutical foundry for space. The first reason is that there are mAb therapies commercially approved or in development for multiple important disease states of spaceflight including osteoporosis^44^, migraines/headaches^45^, seizure^46^, pneumonia^47^, ocular herpes^48^, otitis media^49^, various oncological indications^50^, and fungal infections^51^. A second reason is that a common manufacturing system can be used to produce treatments for a variety of indications which is highly advantageous in mass and volume savings for spaceflight. And thirdly, the economic incentive of research into mAb purification technology has resulted in a plethora of technologies, enabling this analysis to include head-to-head comparisons between multiple mAb capture steps of different origins (e.g., biotic, abiotic) and different processing mechanisms (e.g., bind-and-elute mode liquid chromatography, precipitation). It is in comparing the differences between these technologies that we can uncover general insights into the desired components of a pharmaceutical foundry for space.

Monoclonal antibody therapy is a platform technology that supports human health across a diversity of medical indications with a generally maintained molecular structure, in large part due to the coupling of high target selectivity in the two small and highly variable complementarity-determining regions located in the antigen-binding fragments^52^ and control of the biological action on that target (i.e., effector function) through the generally conserved fragment crystallizable (Fc) region^53^. This otherwise high structural fidelity conserved across mAb therapy products (which are primarily of the immunoglobulin G class) spans a wide variety of therapeutic indications and creates an opportunity for generic mAb production process flows, which include technologies devised specifically for mAb production^54^. This specialized manufacturing, which is most notable in the use of the affinity capture step targeting the Fc region of an antibody with the use of the protein-based ligands derived from the *Staphylococcus aureus* Protein A molecule, can be tuned for highly efficient purification of mAb and antibody-derived (e.g., Fc-fusion protein) class molecules^33^. Other similar protein ligands, such as Protein G and Protein L, are also widely used for their ability to capture different types of immunoglobulin classes and subclasses more efficiently^55^.

#### 2.1.2 Abiotic and biotic Protein A-based unit procedures

We chose to analyze three commercially available abiotic and three development-stage biotic Protein A-based capture step procedures: pre-packed chromatography (CHM), spin column (SPN), magnetic bead (MAG), plant virus-based nanoparticle (VIN), elastin-like polypeptide (ELP), and oilbody-oleosin (OLE) (Figure 1). Commercial technology procedures are based on product handbooks while the procedures of developing technologies are based on reports in literature. This set of procedures was selected to survey a wide range of operational modalities, technological chassis, and perceived advantages and disadvantages (Table 1).

**Figure 1.**
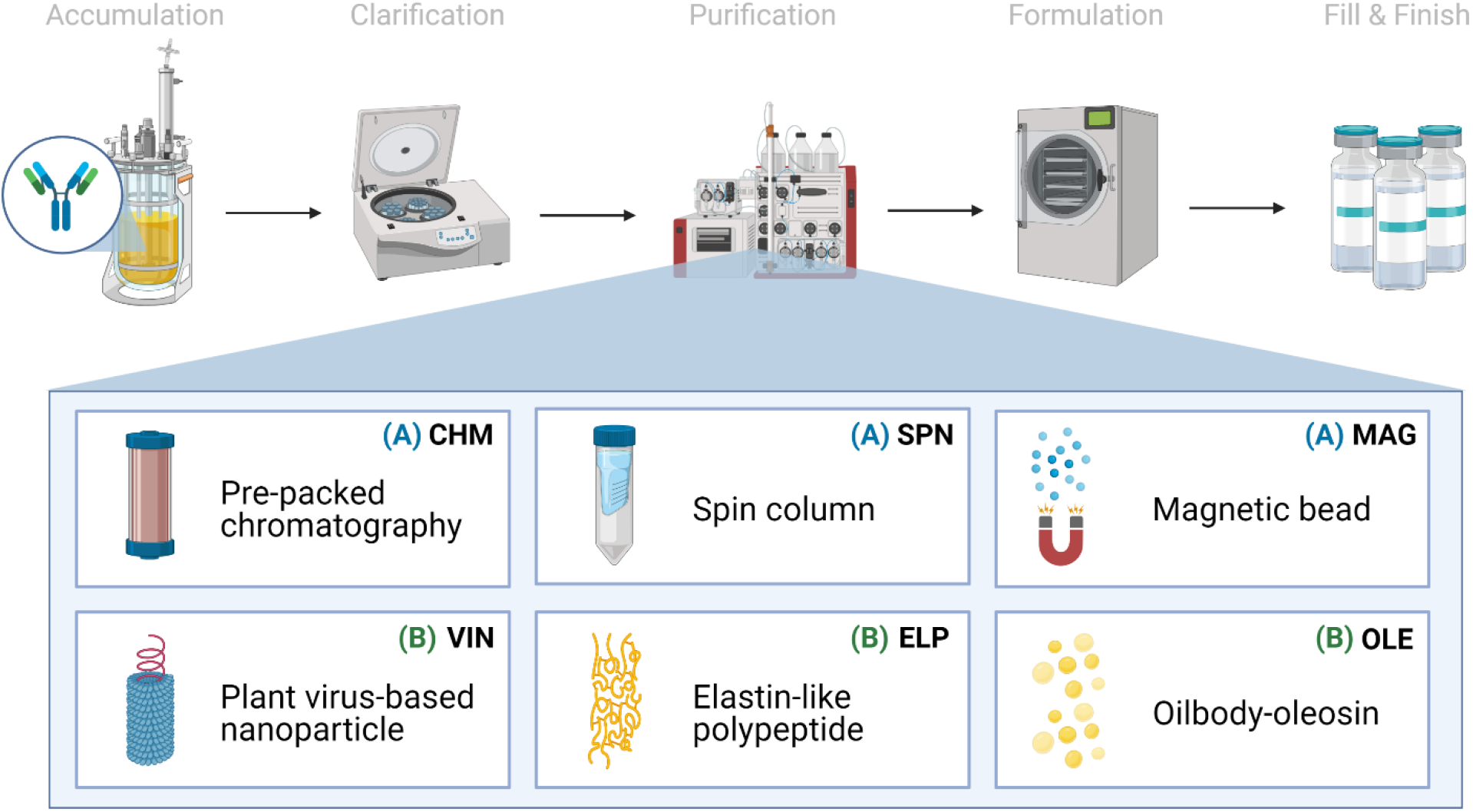
Monoclonal antibody production consists generically of product accumulation, clarification, initial purification, formulation, and fill & finish. Here we investigate six technologies for the capture step within the first purification step in a space mission context using extended equivalent system mass. The manufacturing origin of the capture reagent is denoted as either (A) abiotic or (B) biotic.

**Table 1.** List of Protein A-based monoclonal antibody capture step unit procedures included for analysis.

| Unit Procedure ID | Method | Technology Used | Reference |
| --- | --- | --- | --- |
| Pre-packed chromatography<sup>(A)</sup> | Liquid chromatography | Pre-packed HiTrap MabSelect SuRe column of novel alkali-tolerant recombinant Protein A-based ligand coupled with an agarose matrix | Vendor handbooks<sup>56–58</sup> |
| Spin column<sup>(A)</sup> | Centrifuge-assisted liquid chromatography | Pre-packed Protein A HP SpinTrap spin column containing Protein A Sepharose High Performance |  |
| Magnetic bead<sup>(A)</sup> | Magnetic separation | Protein A Mag Sepharose superparamagnetic beads coupled with native Protein A ligands |  |
| Plant virus-based nanoparticle<sup>(B)</sup> | Sedimentation complex | Plant virion, *Turnip vein clearing virus*, presenting a C-terminal coat protein fusion display of Protein A (domains D & E) | Werner et al. 2006<sup>23</sup> |
| Elastin-like polypeptide<sup>(B)</sup> | Inverse transition cycle | Elastin-like polypeptides (78 pentapeptide (VPGVG) repeats) fused with Z domain, an engineered B domain of Protein A | Sheth et al. 2014<sup>21</sup> |
| Oilbody-oleosin<sup>(B)</sup> | Liquid-liquid partition | *Arabidopsis* oleosin fused at the N-terminal with an engineered Protein A(5) | McLean et al. 2012<sup>59</sup> |

<sup>A</sup>abiotic technology; <sup>B</sup>biotic technology

**Table 2.** Key mission and pharmaceutical reference mission architecture details and assumptions. mAb, monoclonal antibody.

| Mission scope |  |
| --- | --- |
| Pre-deployment | N/A |
| Transit to Mars | 210 days |
| Surface operations | 500 days |
| Return transit | 200 days |
| Total mission duration | 910 days |
| Crew size | 6 crew members |
| Pharmaceutical scope |  |
| Mission demand, mAb | 30,000 mg |
| Biomanufacturing, mAb | 10,000 mg |
| Capture step recovery | 98% |
| Production window | 600 days |
| Feed mAb concentration | 1 mg/mL |
| Molecular weight, mAb | 150 kDa |

All six of the unit procedures are operated in bind-and-elute mode, in which a clarified mAb-containing liquid stream is fed into a capture step containing Protein A-based ligand, which selectively binds the mAb and separates the mAb from the bulk feed stream. The mAb is eluted from the Protein A-based ligand and recovered using a low pH dissociation mechanism. Finally, the low pH environment of the recovered mAb is neutralized for future processing or storage.

CHM is a chromatography system consisting of a liquid sample mobile phase which is pumped through a pre-packed bed of Protein A-fused resin beads housed in a column. SPN is a similar system, in which a Protein A-fused resin bead bed has been pre-packed into a plastic tube housing and the mobile phase flow is controlled via centrifugation of the plastic tube. MAG is a slurry-based magnetic separation system that uses superparamagnetic particles coated with Protein A-fused resin mixed as a slurry with the feed mAb stream for capture and elution of the mAb by magnet. VIN is a sedimentation-based system that uses plant virion-based chassis fused with Protein A-based ligands in suspension for capture of the mAb and centrifugation, assisted by the sedimentation velocity contribution of the chassis, to isolate and elute the mAb. ELP is a precipitation-based system that uses stimuli-responsive biopolymers fused with Protein A-based ligands in suspension for capture of the mAb and external stimuli (e.g., temperature, salt) to precipitate the bound complex and elute the mAb. OLE is a liquid-liquid partitioning system that uses oil phase segregating oleosin proteins fused with Protein A-based ligands to capture mAb in the oil phase and then elute the mAbs into a clean aqueous phase.

### 2.2 Techno-economic evaluation

Techno-economic evaluations are performed using the recently proposed equations for ESM that include calculation of costs at each mission segment^42^. Equivalent system mass (ESM) for the mission 𝔐_0_ is defined as

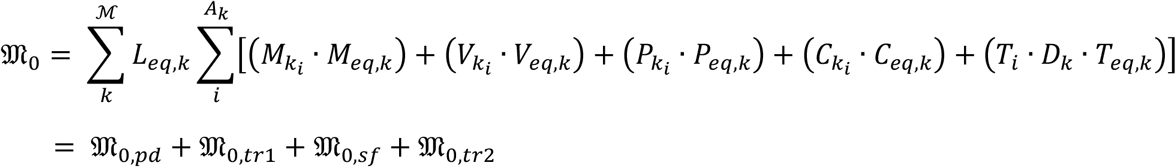

where *M*_*i*_, *V*_*i*_, *P*_*i*_, *C*_*i*_, *T*_*i*_ are the initial mass [kg], volume [m^3^], power requirement [kW], cooling requirement [kg/kW], and crew-time requirement [CM-h/h], *M*_*eq*_, *V*_*eq*_, *P*_*eq*_, *C*_*eq*_, *T*_*eq*_ are the equivalency factors for mass [kg/kg], volume [kg/m^3^], power [kg/kW], cooling [kg/kW], and crew time [kg/CM-h], respectively, *L*_*eq*_ is the location equivalency factor [kg/kg], and *D* is the duration of the mission segment [day] over a set of subsystems *i* ∈ *A* and set of mission segments *k* ∈ ℳ. The mission ESM in this study is specifically defined as the sum of subtotal ESM for each mission segment within the scope of the reference mission architecture defined in this study (pre-deployment 𝔐_0,*pd*_, crewed transit to Mars 𝔐_0,*tr*1_, Mars surface operations 𝔐_0,*sf*_, and return crewed transit to Earth 𝔐_0,*tr*2_).

The mission timeline depicted in Figure 2 provides insight into the proposed RMA and downstream crew needs and mAb production horizon. Here we assume a total mission duration of 910 days. First, a crew of 6 will travel from Earth to low Earth orbit, then board an interplanetary craft for a 210-day journey to Martian orbit, where the crew will descend to the surface in a separate craft, allowing the large transit vehicle to remain in orbit. Once on Mars, the crew will perform surface operations for 600 days. Following surface operations, the crew will leave Mars in a fueled ascent craft, board the interplanetary vehicle, and return to Earth orbit in 200 days.

**Figure 2.**
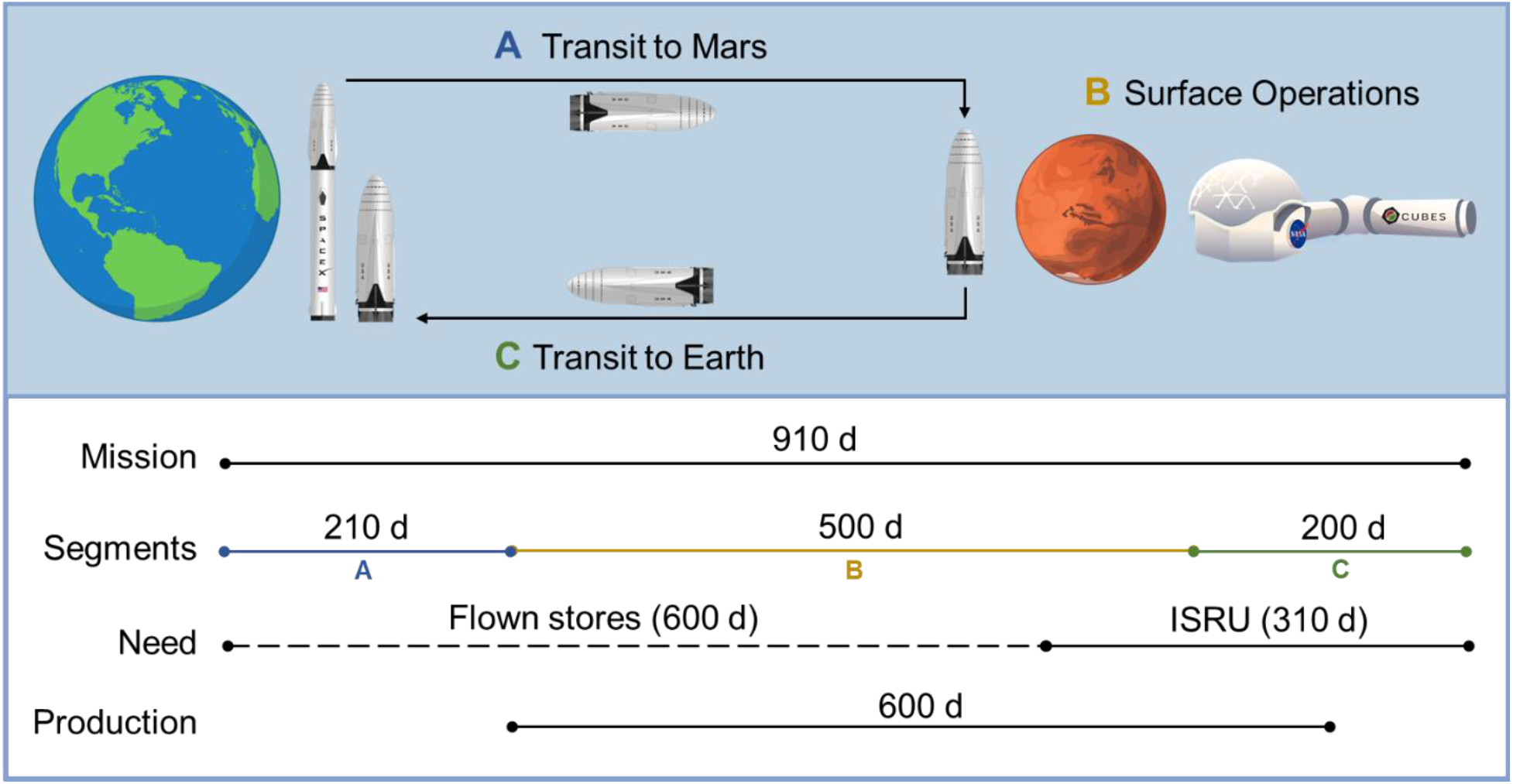
An illustration of the reference mission architecture in which (A) a crewed ship is launched from the surface of Earth and lands on Mars and (B) assembles a pre-deployed habitat on the Martian surface to perform operations before (C) a return transit to Earth on the same ship. Pharmaceutical needs are supported by flown stores until partway through surface operations, at which point needs are met by pharmaceuticals produced using *in situ* resource utilization. Production is initiated prior to the need window to ensure adequate stocks are generated by the time it is needed. Rocket artwork adapted from Musk, 2017^60^. Habitat artwork by Davian Ho.

The mission demand for mAb therapies is assumed to be 30,000 mg over the entirety of the mission (supporting logic detailed in Supplementary Information, Table S1). Pharmaceutical stores and production resources are assumed to be flown with the crew transit (no pre-deployment in order to maximize shelf-life). It is assumed that the first 600 days of pharmaceutical demand will be met through flown stores (20,000 mg), at which point pharmaceutical ISRU manufacturing is needed (10,000 mg) to alleviate the impact of accelerated pharmaceutical degradation and provide supplementary medication. The pharmaceutical production window opens prior to the ISRU demand timeframe and persists through a portion of the return transit (up to mission day 810) to reflect the expected life support advantage of maintaining capabilities to counter unanticipated needs or threats. We assume that the Protein A-based unit procedures consistently yield 98% recovery of mAb from the input stream.

### 2.3 Unit procedure simulation

Deterministic models for each unit procedure were developed in Microsoft Excel (Supplementary Information, Spreadsheet S1) using reference protocols cited in Table 1 as a series of executable operations, each containing a set of inputs defined by cost categories (labor, equipment, raw materials, consumables) that are correspondingly populated with characteristic ESM constituent (mass, volume, power, cooling, labor time) values (model composition illustrated in Supplementary Information, Figure S1). Unit procedures have been defined as the smallest single execution (i.e., unit) of the secondary purification capture step procedure according to the reference protocol. We define the unit capacity by volume according to the equipment and consumables used (e.g., 2 mL maximum working volume in a 2 mL tube) and by mAb quantity according to the binding capacity for the given method (e.g., 1 mg mAb/mL resin) (Supplementary Information, Table S2). Unit procedures with no explicit working volume constraints (i.e., the liquid solution volume for biotic technologies) have been defined with a maximum unit volume of 2 mL. ESM-relevant characteristics of individual inputs (e.g., equilibration buffer, 2 mL tube) are defined based on publicly available values, direct measurements taken, and assumptions (which are explicitly identified in the spreadsheet).

There are several model features that we have considered and decided not to include within the scope of analysis. Packing and containers for the inputs are not included for three reasons: 1) the contributions of the container are considered negligible as compared to the input itself (e.g., container holding 1 L buffer as compared to the 1 L of liquid buffer); 2) materials flown to space are often re-packaged with special considerations^61^; and 3) the selection of optimal container size is non-trivial and may risk obscuring more relevant ESM findings if not chosen carefully. We do not consider buffer preparation and assume the use of flown ready-to-use buffers and solutions. Furthermore, refrigeration costs of the input materials and costs that may be associated with establishing and maintaining a sterile operating environment (e.g., biosafety cabinet, 70% ethanol in spray bottles) are expected to be comparable between unit procedures and not considered. Refrigeration costs associated with low temperature equipment operation (e.g., centrifugation at 4 °C) are included in the equipment power costs.

Inputs common across unit procedures are standardized (Supplementary Information, Table S3). One operational standardization is the inclusion of pH neutralization of the product stream following the low pH elution mechanism, which was explicitly stated in some procedures while not in others. Input quantities are scaled from a single unit to determine the number of units required to meet the reference mission architecture specifications. The ESM constituent inputs (mass, volume, power, cooling, labor time) are converted into equivalent mass values using the RMA equivalency factors (Supplementary Information, Equation S1 & Table S4).

## 3. Results & Discussion

### 3.1 Standardization of manufacturing efficiency

Given the limited granularity of the presented reference mission architecture, which was scoped as such to reflect the lack of literature presenting an overarching and validated Concept of Operations for a Transit to Mars^62^, we do not define strict manufacturing scheduling criteria for pharmaceutical production. Construction of a detailed pharmaceutical production RMA is hindered by uncertainty in the number and identity of mAb therapy products that would be included within mission scope, the decay rate of mAb therapy stores in the mission environments, and a reasonable basis for building robustness to unanticipated disease states. Rather, we choose to establish an objective comparison between unit procedures by normalizing for scheduling-associated manufacturing efficiencies. We accomplish this by first identifying the number of batches per mission (and thus batch size) needed to meet the mAb demand (base case of 10,204 mg mAb feed assuming 98% recovery) that minimizes the ESM output for a given unit procedure, and then running the simulation of pharmaceutical production at that number of mission batches, as shown in Figure 3a and tabulated in Supplementary Information, Table S5.

**Figure 3.**
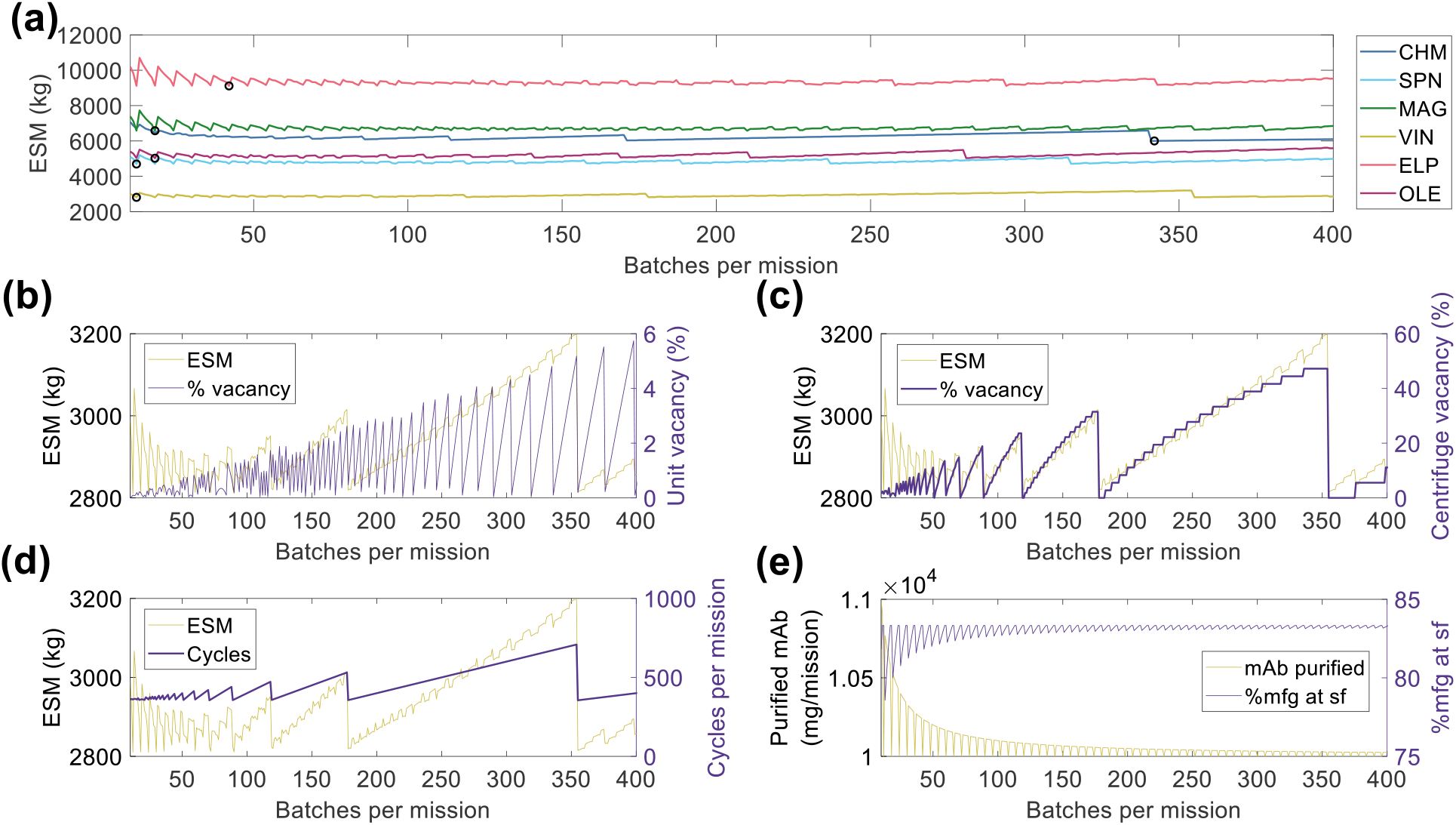
(a) Scheduling optimization for the establishment of base case scenarios for each unit procedure. The value for number of batches corresponding to the minimum equivalent system mass for each unit procedure, as indicated by black circle (**○**) markers. Key operational parameters impacted by mission scheduling (shown using the VIN procedure) include (b) unit underutilization or vacancy, (c) equipment underutilization or vacancy, in this case represented by the centrifuge as the bottleneck, (d) the number of use cycles, and (e) the total quantity of monoclonal antibody (mAb) per mission and per surface operation (sf). CHM, pre-packed chromatography; SPN, spin column; MAG, magnetic bead; VIN, plant virus-based nanoparticle; ELP, elastin-like polypeptide; OLE, oilbody-oleosin.

In Figure 3b – e, we visualize a deconstruction of ESM output, using the VIN unit procedure as an example, by key performance metrics that vary with a scheduling dependence in order to illustrate the significance of batch optimization in unit procedure comparison. The processing of a given batch volume and mAb quantity is allocated into a number of units, as determined by the volume and mAb quantity constraints of a given unit procedure, and a number of use cycles per batch, as determined by the capacity of the equipment specified in the given unit procedure. We show how the variation in ESM output over the number of mission batches maps to extent of unit vacancy or underutilization (Figure 3b), extent of operational equipment (e.g., centrifuge) vacancy or underutilization (Figure 3c), and number of required use cycles (Figure 3d). We also show an oscillatory behavior in the scheduling (i.e., total mAb purified per mission, % purified at surface operations) that quickly dampens as number of mission batches increases (Figure 3e). This behavior is a result of the assumption that the mAb feed stream is coming from a discrete upstream production batch (e.g., batch-mode bioreactor) that does not output partial batch quantities, as opposed to a continuous upstream production for which there are no defined batches. Accordingly, partial batch needs are met by the processing of a full batch.

### 3.2 Base case scenario

The ESM and output metrics of the base case scenario (10,000 mg mAb demand, 1 mg mAb/mL feed concentration, 98% recovery) for each of the six unit procedures are shown in Figure 4a-f. From this viewpoint of an ESM output for an isolated unit procedure outside the context of a full purification scheme, the ESM ranked from lowest to highest are VIN < SPN < OLE < CHM < MAG < OLE. However, we reason that it is more important to understand the model inputs that influence the ESM output rankings than to use the rankings in this isolated subsystem analysis to make technology selection choices, which requires the context of a full pharmaceutical foundry and of linkages to other mission elements.

**Figure 4.**
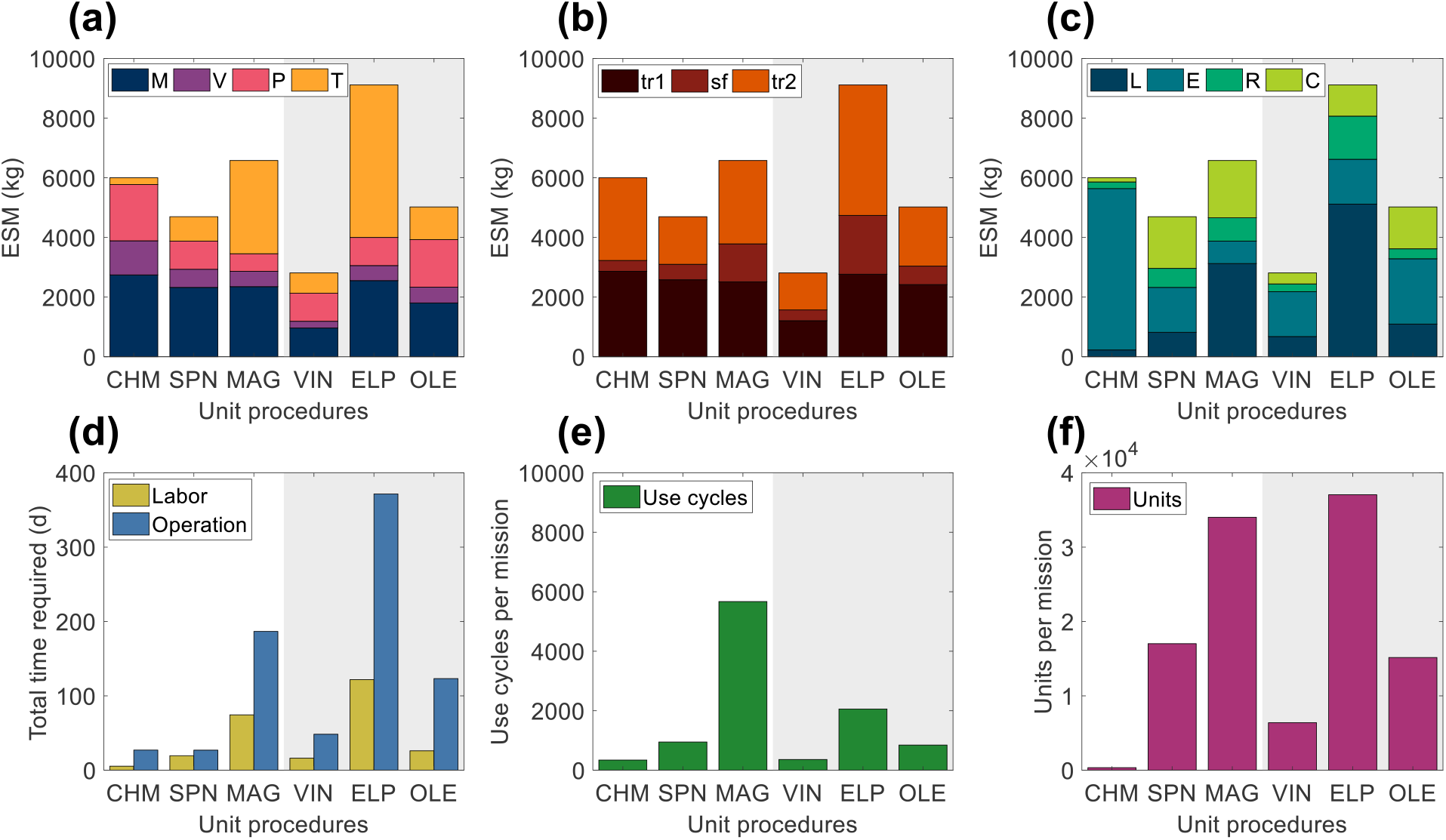
Base case equivalent system mass results broken down by (a) mass (M), volume (V), power (P), and labor time (T) constituents, (b) transit to Mars (tr_1_), surface operations (sf), and return transit (tr_2_) mission segments, and (c) labor (L), equipment (E), raw materials (R), and consumables (C) cost category for the six tested Protein A-based monoclonal antibody affinity capture step unit procedures segregated by abiotic (white background) and biotic (grey background) technologies. Also shown are the (d) labor and operation times, (e) number of use cycles, and (f) number of units required for each unit procedure to meet the reference mission demand. CHM, pre-packed chromatography; SPN, spin column; MAG, magnetic bead; VIN, plant virus-based nanoparticle; ELP, elastin-like polypeptide; OLE, oilbody-oleosin.

We observe that mass costs are generally the primary contributor to ESM output, except for the MAG and ELP procedures in which labor time costs are larger. The mass costs are not closely associated to any given cost category across unit procedures, but rather the breakdown of mass costs varies widely by unit procedure.

Power costs (kW) are disproportionately high given that the static nature of ESM assumes constant usage, and thus energy (kW·h) in this context (i.e., the power supply to the equipment is not turned off in this analysis). Time of power useage as a fraction of duration are as follows: CHM (99%) > MAG (78%) > ELP (48%) > SPN (45%) > OLE (42%) > VIN (30%). The electrical needs of the equipment used by the unit procedures are within NASA-proposed Mars mission RMA bounds, with energy use across all unit procedures would peak at ∼1% of a proposed Mars transfer vehicle electric capacity (50 kWe) or ∼5% of the habitat capacity (12 kWe) of a reference stationary surface nuclear fission power reactor^63^.

The mission segment breakdown of ESM illustrates the relatively high costs of pharmaceutical manufacturing capabilities for transit, even for the transit to Mars (tr_1_) in which there is no actual production taking place. There is a strong economic incentive to limit the amount of supplies flown on tr_1_ . Alternatives such as the pre-deployment of reagents and consumables and limiting of production to surface operations on Mars (which has lower RMA equivalency factors for mass and volume than transit operations) must be balanced against the risk to human health posed by removing pharmaceutical production capabilities from a mission segment and potentially exposing the supplies to longer storage times that could challenge shelf lives.

Labor and operation times are important parameters in the broader mission and pharmaceutical foundry context. These unit procedures represent a single step of pharmaceutical production, which if realized in a space mission context, would, in turn, need to be a small portion of a crew member’s time allocation. Assuming 40-hour work weeks for crew members, the labor time spans a range of ∼1% (CHM) to ∼14% (ELP) of the available crew time over the 600-day production window. It is not feasible to operationalize with such high labor and operation times at this scale of production, particularly as they stand for MAG and ELP. While strategies such as batch staggering and concurrency can be used to reduce durations, advanced automation will almost certainly need to be built into the core of a pharmaceutical foundry.

A prevailing trend throughout the unit procedures is that the number of unit executions and use cycles required by a given unit procedure are positive correlated with the ESM output value, except for the equipment cost-dominant and higher unit capacity CHM procedure. The equipment modeled in the analysis for CHM and the other unit procedures are almost certainly not space-ready and could be further designed to reduce mass and volume and increase automation to reduce crew labor time. The increased equipment costs in the CHM procedure are primarily due to automation and monitoring hardware for running liquid chromatography, which is reflected in the minimal labor costs of the CHM procedure. Miniaturization efforts, such as those focusing on microfluidic systems^64–66^, are emerging as a potential path towards mitigating the high equipment costs associated with highly automated and tightly controlled manufacturing, which are crucial for freeing up valuable crew time.

The number of unit executions is determined by the binding capacity of the technology and the nominal unit size. This indicates that the unit capacity for purification is an important consideration and influential factor. Unit sizing is an important consideration that is valuable to assess more holistically within the broader pharmaceutical production and mission context.

The number of use cycles is determined by the number of unit executions required and by the maximal unit capacity of the equipment items (e.g., if you presume that an 18-slot centrifuge is the equipment bottleneck then the effective number of batches is the number of units required divided by 18). Therefore, it can be understood that the equipment unit capacity is a critical parameter in tuning the number of use cycles and, by extension, the labor costs. For processes with lower labor costs, due to the intrinsic nature of the procedure or through automation of labor, equipment unit capacity will still influence the total duration and production throughout. The MAG and ELP procedures yield both high labor and duration times and are thus particularly sensitive to the equipment capacity.

To contextualize the ESM results of this analysis, we compare them to recently reported ESM outputs of a Mars surface greenhouse^67^. In this study, they report an ESM output of ∼200 – 1,200 kg per kg plant biomass, excluding consumable costs (i.e., plant growth substrate, nutrient salts, cleaning agents), which they term as logistics mass. For proper comparison to these values, we removed mission segment dependencies to evaluate only surface operations ESM and changed equivalency factors to those used in the Mars surface greenhouse study (Supplementary Information, Table S6). With these modifications in place, we calculate ESM totals of ∼300 – 1,700 kg per 10,000 mg mAb. From the perspective that a crewed Mars surface mission is projected to require ∼10,000 kg food mass^32^, and that the studied crop harvest indices varying from 40 – 90%^31^, it frames the ESM for this portion of pharmaceutical purification as a very minor fraction of food life support costs.

#### 3.2.1 Contextualizing ESM with supporting evaluations

Having acknowledged shortcomings of ESM as a decision-making tool for comparison of alternative approaches in isolated subsystems, we propose that supplementary evaluations can assist in contextualization. A primary gap of an isolated subsystem ESM analysis is a lack of information on the holistic usefulness or cost of a given employed resource, which could include its synergy with other mission subsystems and its extent of recyclability, or waste loop closure, within the mission context. For example, the isolated subsystem analysis does not capture information on the broad applicability that a centrifuge might have for use in other scientific endeavors, nor do the ESM outputs reflect the > 93% recyclability of water achieved by the recycler on the ISS^68^ that may be generalizable to future missions.

The use of environmental footprint metrics, such as PMI, may be one valuable step towards capturing missed information on recyclability. PMI a simple metric of material efficiency defined as the mass of raw materials and consumables required to produce 1 kg of active pharmaceutical ingredient. The study by Buzinski et al. introducing PMI for biopharmaceuticals presents data from 6 firms using small-scale (2,000 – 5,000 L reactor) and large-scale (12,000 – 20,000 L reactor) mAb manufacturing operations, finding an average 7,700 kg of input is required to produce 1 kg of mAb^36^. Figure 5a presents PMI evaluation for the six capture steps included in analysis, which result in PMI outputs as low as 2,390 kg of input (CHM) and as high as 17,450 kg of input (MAG) per 1 kg of mAb. A comparison of these outputs to those of Budzinski et al. indicates that we may be observing roughly similar values after accounting for the high cost of initial purification in the study, representing ∼60% of the total PMI reported, the elevated feed mAb concentration (i.e., cell culture titer) of 1 – 5.5 g mAb/L, and adjustments for economies of scale when operating at such low cycle volumes (Figure 5b). Consumable costs appear to be the most sensitive to scale, which represents ∼1% total PMI on average in the values reported by Budzinski et al. and ranges from 35% (CHM) to 77% (OLE) here. Budzinski et al. also go one step further to distinguish water as a separate category from raw materials and report that >90% of the mass is due to water use. Here we assume pre-made buffers and do not directly add water in this study, so we refrain from a similar calculation, but it is worth noting that the extent of water use may also serve as a reasonable starting surrogate for extent of achievable recyclability in a space mission context.

**Figure 5.**
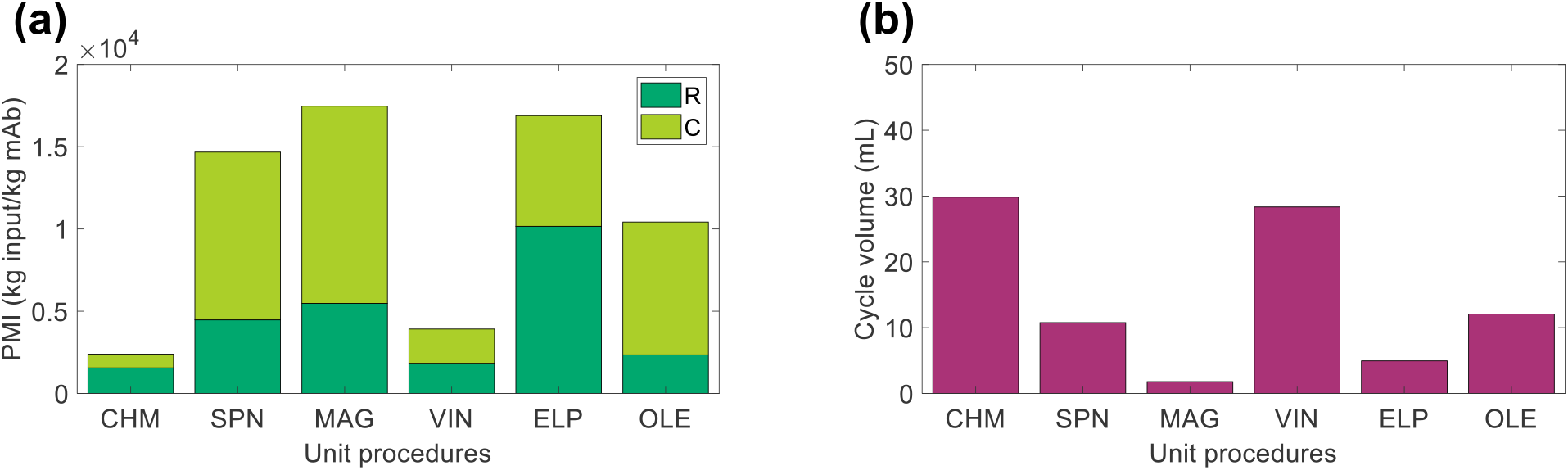
Process mass intensity (PMI) evaluation of the unit procedures broken down by raw materials (R) and consumables (C) contributions. CHM, pre-packed chromatography; SPN, spin column; MAG, magnetic bead; VIN, plant virus-based nanoparticle; ELP, elastin-like polypeptide; OLE, oilbody-oleosin.

### 3.3 Scenario analysis

We analyzed the specific ESM output broken down by cost category for the six unit procedures over a range of input stream mAb concentrations (Figure 6) and mission demand for mAb (Figure 7). Specific ESM, termed cost of goods sold in traditional manufacturing analyses, is the ESM output required to produce 1 mg mAb. This is used in the scenario analyses to normalize ESM output across variation in mission demand for mAb. The optimal number of batches per mission was found and used for each unit procedure and scenario tested (Supplementary Information, Tables S7 – S8).

**Figure 6.**
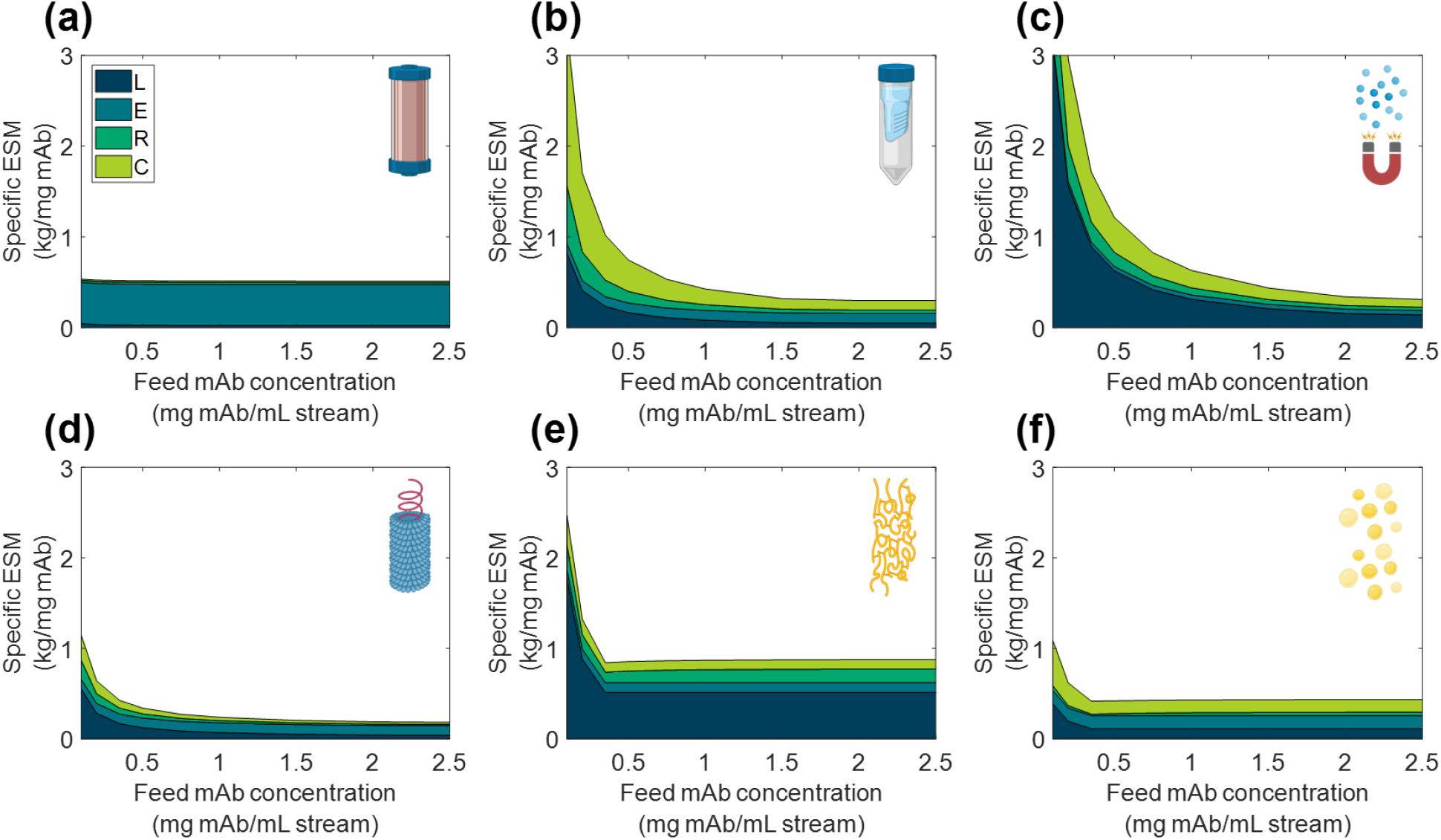
Specific equivalent system mass (per unit mass monoclonal antibody produced) broken down by labor (L), equipment (E), raw materials (R), and consumables (C) cost categories as a function of feed monoclonal antibody (mAb) concentration for (a) CHM, (b) SPN, (c) MAG, (d) VIN, (e) ELP, and (f) OLE. CHM, pre-packed chromatography; SPN, spin column; MAG, magnetic bead; VIN, plant virus-based nanoparticle; ELP, elastin-like polypeptide; OLE, oilbody-oleosin.

**Figure 7.**
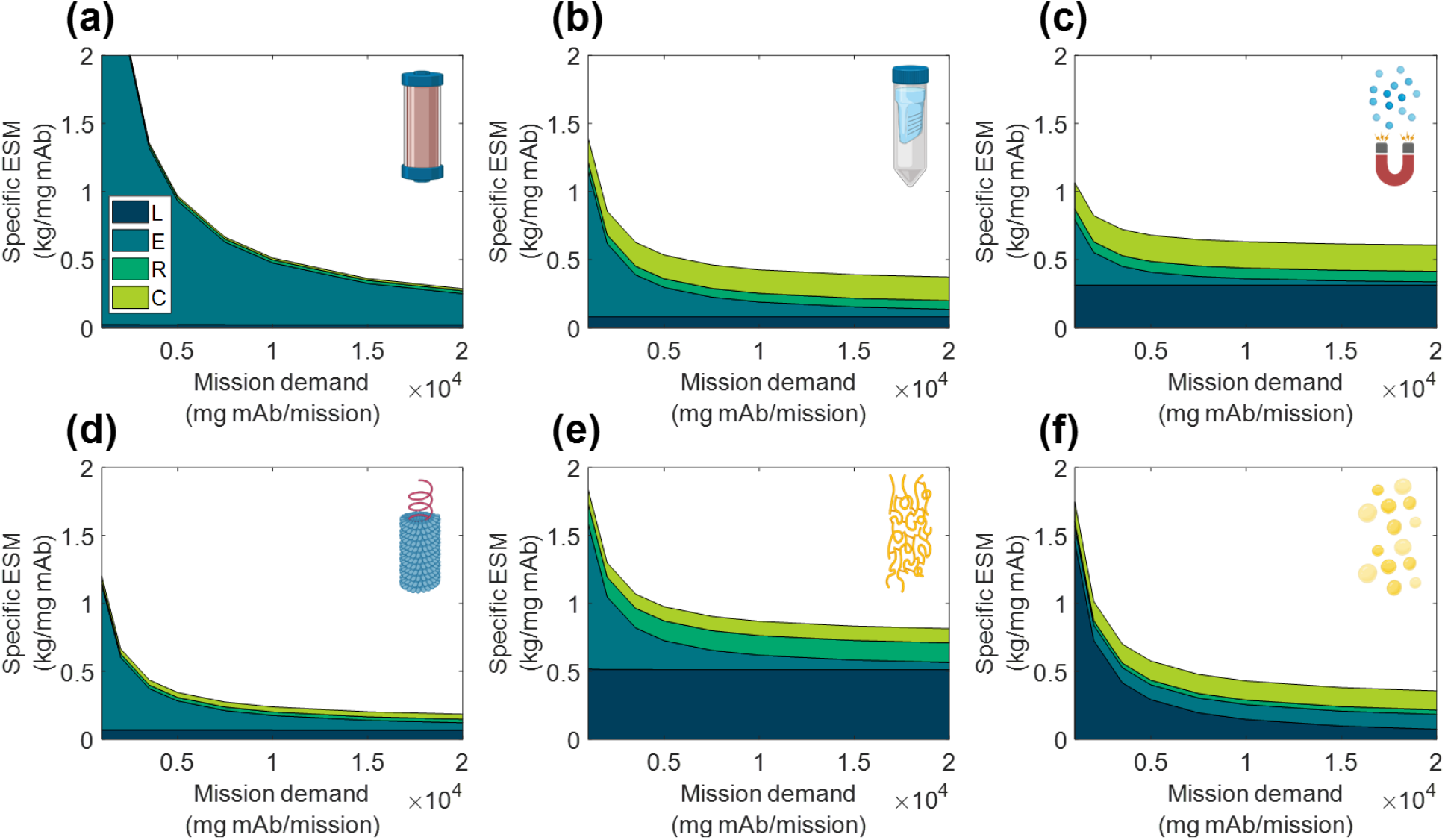
Specific equivalent system mass (per unit mass monoclonal antibody produced) broken down by labor (L), equipment (E), raw materials (R), and consumables (C) cost categories as a function of mission production demand for monoclonal antibody for (a) CHM, (b) SPN, (c) MAG, (d) VIN, (e) ELP, and (f) OLE. CHM, pre-packed chromatography; SPN, spin column; MAG, magnetic bead; VIN, plant virus-based nanoparticle; ELP, elastin-like polypeptide; OLE, oilbody-oleosin.

We observe the general and expected trends that specific ESM decreases with an increasing feed stream mAb concentration and mission demand. The CHM procedure exhibits notably limited sensitivity to feed stream mAb concentration, which can be attributed to the equipment-dominated cost profile, fixed column size, and nature of the governing reference protocol that does not specify restrictions on sample load volume. Depending on the pre-treatment of the feed stream, it may be more reasonable to impose constraints on the sample load volume. In contrast, the specific ESM output of the CHM procedure is the most sensitive to mission mAb demand with higher demand increasingly offsetting the fixed capital costs. The CHM procedure is also the largest capacity unit modeled in the analysis (i.e., CHM capacity is 30 mg mAb/unit as compared to 2.7 mg mAb/unit for MAG, the next highest capacity unit) and is accordingly expected to scale well with demand.

The SPN, ELP, OLE procedures exhibit behaviors in which the specific ESM output abruptly plateaus with an increasing feed stream mAb concentration. This observation can be attributed to the unit procedure operating in a mAb binding capacity-limited regime (as opposed to volume-limited for more dilute feeds) which also then controls and maintains unit procedure throughput (e.g., the ELP number of units, 37,044, and use cycles per mission, 2,058, is constant at and above 0.35 mg mAb/mL input stream concentration). This can be de-bottlenecked via technology (e.g., improved chemistry of the capture step unit leading to higher binding capacity) or methodology (e.g., increased concentration of the capture step unit leading to higher binding capacity) improvements.

Low demand scenarios are particularly relevant for examination in a space health context, as small capacity redundant and emergency utility is a likely proving ground for inclusion of a space pharmaceutical foundry. At the lower boundary of the tested range (1,000 mg mAb/mission), we see the ESM outputs from lowest to highest are re-ordered as MAG < VIN < SPN < OLE < ELP < CHM. Minimization of equipment costs are particularly important in this regime, and it is observed that, indeed, the ESM output near completely aligned with the ranking of equipment cost (MAG < VIN < SPN < ELP < OLE < CHM). It is likely that other non-ESM factors such as integration with other flown elements will understandably influence the design and composition of early and low capacity flown pharmaceutical foundries.

### 3.4 Alternate scenarios

#### 3.4.1 Mission configurations

We explored variations to the base case RMA for all six unit procedures including scenarios in which the pharmaceutical manufacturing resources are shipped prior to the crew in pre-deployment, (+)pd, the production window has been truncated to close with the end of surface operations, (-)tr_2_, and a combination of the two prior modifications, (+)pd (-)tr_2_ (Figure 8). Costs of pre-deployment are included in the analyses and mission demand is kept constant regardless of the production window.

**Figure 8.**
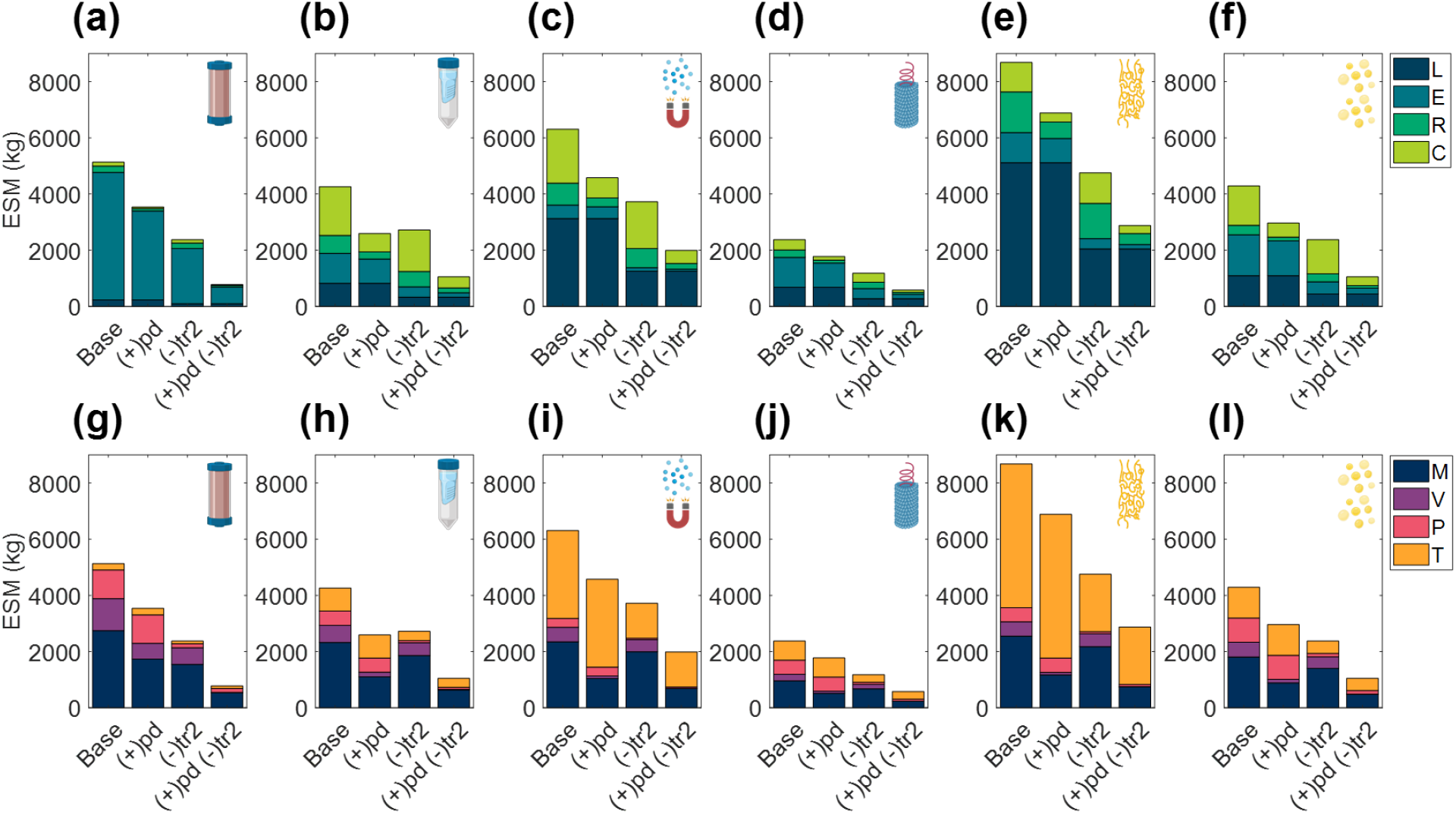
Evaluation of extended equivalent system mass values in various mission configurations broken down by labor (L), equipment (E), raw materials (R), and consumables (C) cost categories cost category and mass (M), volume (V), power (P), and labor time (T) constituents for CHM, (a, g), SPN, (b, h), MAG, (c, i), VIN, (d, j), ELP (e, k), and OLE, (f, l). Configurations include the base case scenario of manufacturing resources flown with the crew for pharmaceutical production on the surface and return transit (Base), and alternatives in which the manufacturing resources are flown prior to the crew in pre-deployment, (+)pd, the production window is limited to surface operations, (-)tr_2_, and a combination of the two previously stated alternatives, (+)pd (-)tr_2_ . CHM, pre-packed chromatography; SPN, spin column; MAG, magnetic bead; VIN, plant virus-based nanoparticle; ELP, elastin-like polypeptide; OLE, oilbody-oleosin.

In all cases the ESM totals were reduced from the base case. Additionally, the general trend held that (-)tr_2_ scenario resulted in lower ESM totals than (+)pd scenario except for SPN, in which the increased raw material and consumable costs of (-)tr_2_ were sufficiently large to outweigh the reduction in equipment and labor costs of (+)pd. The combination (+)pd (-)tr_2_ scenario resulted in the lowest ESM totals at a fraction of the base case (as high as 39% reduction in SPN and as low as 21% reduction in ELP).

#### 3.4.2 Equipment & unit throughput

Acknowledging the significance of the equipment capacity on ESM output, we further explored this contribution by comparing the base case ESM output of the centrifuge-utilizing procedures (SPN, VIN, ELP, OLE) to that resulting from the use of alternative centrifuge models (Supplementary Information, Table S9). This effectively results in a trade of equipment costs and batch throughput. The optimal number of batches per mission was found and used for each unit procedure and interval tested (Supplementary Information, Table S10).

We observe in Figure 9 that the ESM values increased with the size of the centrifuge model, 12-slot < 18-slot (base) < 48-slot. The labor and consumables savings of higher batch throughput were outweighed by the higher equipment costs (including higher power costs). Operation duration is an important metric relevant to a pharmaceutical foundry that is not well reflected in ESM that is also impacted by this alternative scenario. The exception to this trend is the 48-slot condition for the ELP procedure, in which a lower consumable cost related to the number of use cycles per mission (i.e., pipette tips, tubes, gloves) sufficiently lowered the total ESM below the 18-slot condition.

**Figure 9.**
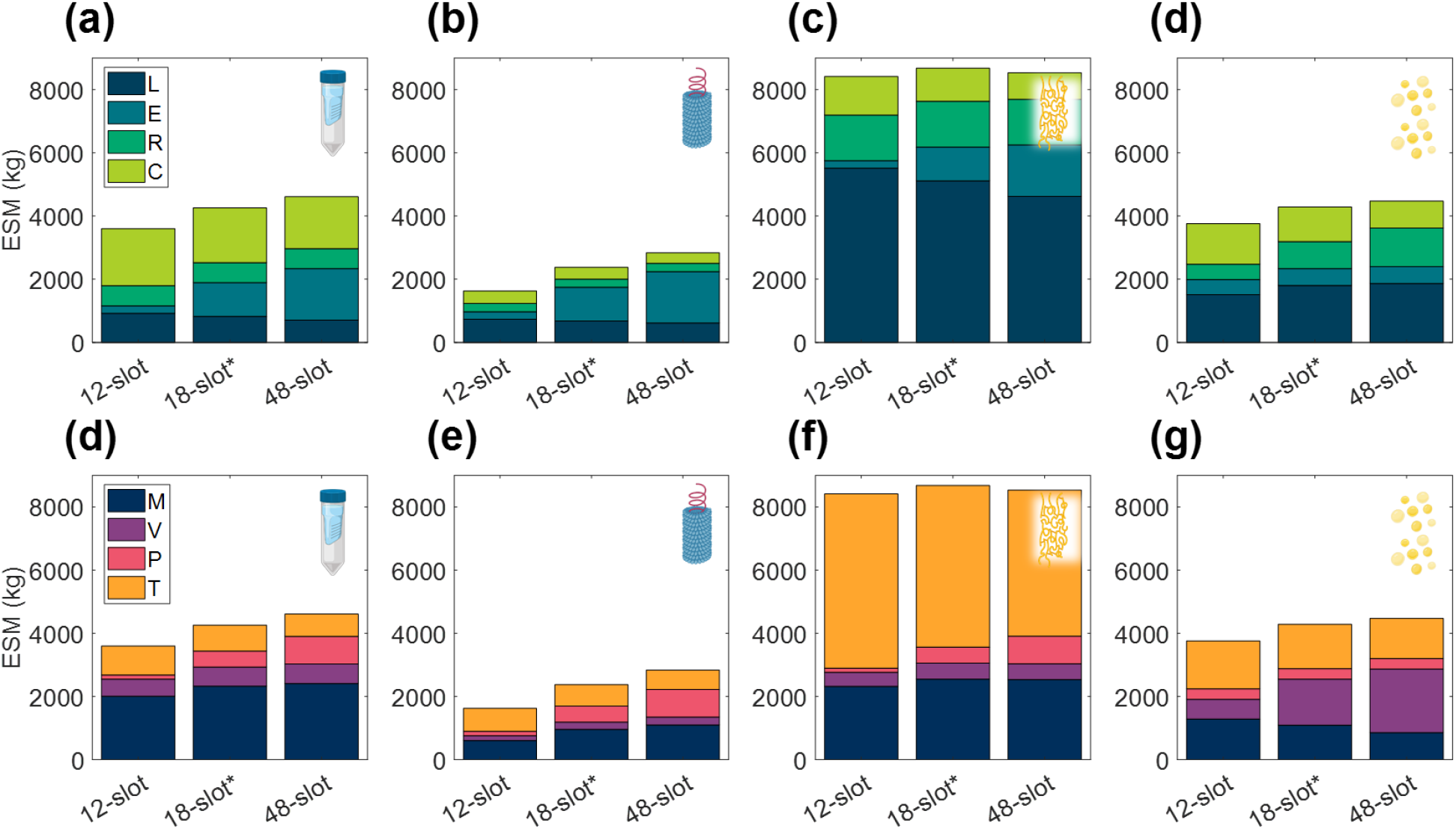
Changes in extended equivalent system mass values with different capacity centrifuge models broken down by labor (L), equipment (E), raw materials (R), and consumables (C) cost categories and mass (M), volume (V), power (P), and labor time (T) constituents for SPN (a, d), VIN (b, e), ELP (c, f), and OLE (d, g). SPN, spin column; VIN, plant virus-based nanoparticle; ELP, elastin-like polypeptide; OLE, oilbody-oleosin.

#### 3.4.3 Technology reusability

The number of use cycles for liquid chromatography resins is an important economic parameter in commercial pharmaceutical manufacturing^71^. Here we explore the impact of use cycles on the CHM and ELP procedures in a space mission context, looking at no reuse nor regeneration operation of the purification technology, (-)Reuse, and at an increased number of use cycles, (+)Reuse (Figure 10).

**Figure 10.**
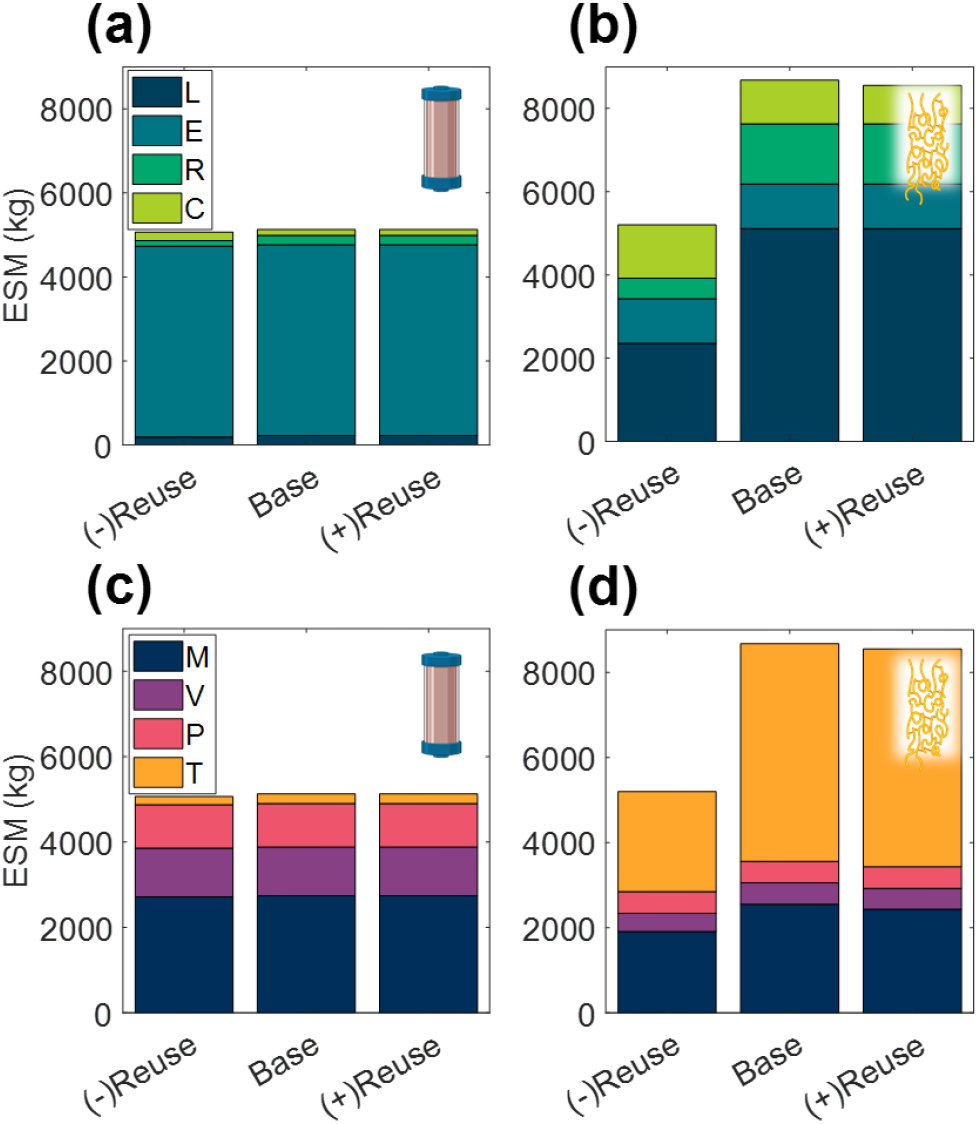
Changes in extended equivalent system mass values with reusability of purification technology broken down by labor (L), equipment (E), raw materials (R), and consumables (C) cost categories and mass (M), volume (V), power (P), and labor time (T) constituents for CHM (a, c), and ELP (b, d). (-)Reuse considers the technology as single-use and accordingly discards the unit procedure cleaning operations; (+) Reuse considers additional reuse cycles of the technology. CHM, pre-packed chromatography; ELP, elastin-like polypeptide.

We observe that the terrestrial importance of use cycles does not prevail in this isolated ESM evaluation in a space context. The high purchase costs of resin are not considered in ESM and the impact of the reuse cycles is reduced to the mass and volume savings of the pre-packed column consumable. There is a minor decrease in ESM of the (+)Reuse over the base case scenario, but both of these result in substantially higher ESM than the (-)Reuse scenario, particularly for the ELP procedure, in which the regeneration operation has been removed in addition to the reusability of the technology.

These results echo the trend of single-use technology in commercial biotechnology in which manufacturers look to disposable plastic bioreactor and buffer bags as a means to reduce cleaning and validation costs^72^. It would be valuable to further consider the utilization of single-use technology in a space pharmaceutical foundry, and in other space systems bioengineering applications, but it is important to point out the limited scope of this ESM analysis. Here we reiterate that the single unit procedure scope establishes a modular basis for pharmaceutical foundry ESM evaluation but does not realize the true circular economy advantages of reuse, which may be considerable for the regeneration step, and of biological systems for production of the purification reagent in general.

## 4. Conclusion & Future Directions

In this study, we have introduced and rigorously applied the ESM framework to biopharmaceutical processing as a first step towards modeling and understanding the costs of Space Systems Bioengineering and, more specifically, of a long-duration space exploration medical foundry, which we believe may one day constitute a critical bioregenerative component of Environmental Control and Life Support Systems for humans to be able to explore the surface of Mars. We have observed that the static behavior of ESM, while certainly maintaining usefulness in early-stage analyses, may stymie later-stage analyses of bioregenerative life support technologies, which tend to behavior more dynamically than traditional abiotic counterparts. This limitation is particularly noticeable in the static ESM power requirements, which would be greatly improved in considering energy dynamics for proper accountancy and integration with specifications of mission electric capacity.

The encouraging comparison of ESM penalties to that of a Mars surface greenhouse motivates future investigation into the ESM output of a complete pharmaceutical foundry for a more complete comparison to other life support system needs and subsequent formal evaluations of medical risk (i.e., loss of crew life, medical evacuation, crew health index, risk of radiation exposure-induced death from cancer) mitigation as a balance to the ESM costs. The Integrated Scalable Cyto-Technology system^19^, reported in literature as capable of “end-to-end production of hundreds to thousands of doses of clinical-quality protein biologics in about 3 d[ays],” is an automated and multiproduct pharmaceutical manufacturing system that may serve well as a starting point for a complete pharmaceutical foundry evaluation.

Assembly of a complete pharmaceutical foundry ESM model would also enable investigation of more nuanced RMA design considerations, such as those relating to the influence of a fixed set, or anticipated probability distribution, of pharmaceutical product diversity and batch size on optimal system composition to meet given medical risk thresholds.

As stated in the original presentation of ESM theory and application, comparison of multiple approaches for a given subsystem with ESM, such as we are studying with the capture step a mAb pharmaceutical foundry, should satisfy the same product quantity, product quality, reliability, and safety requirements^69^. Of these assumptions, the product quality and safety requirements prove challenging for implementation in pharmaceutical foundry comparisons. It is worth noting that reliability is not considered in the scope of this preliminary study, given the varying technology readiness levels of the unit procedures, but that it should be included in future analyses of full purification schemes. High product sensitivity to process changes, and the large battery of testing sometimes required to observe them (the extent of which will also change with the processes employed), creates a situation where ESM comparisons of pharmaceutical foundries that serve as technology decision making tools will absolutely need to meet this requirement, albeit at a considerable cost and/or complexity of execution.

The assessment of equivalent safety requirements, to the best of the knowledge of the authors, has been approached thus far in an ad hoc and qualitative manner, relying on extensive subject manner expertise and working process knowledge. One promising route to strengthening these critically important safety assessments would be to implement a formal assessment framework based on the environmental, health, and safety (EHS) assessment proposed by Biwer and Heinzle^70^, in which process inputs/outputs are ranked based on a series of hazard impact categories (e.g., acute toxicity, raw material availability, global warming potential) and impact groups (e.g., resources, organism). The key to a systematic space health-centric safety assessment like this is to establish space-relevant EHS impact categories (e.g., planetary protection, crew and ship safety). An improvement of the EHS underpinnings has the potential to provide significant benefits to future ESM analyses in the increasingly complex mission architecture of longer-duration missions.

## 5. Acknowledgments

This material is based upon work supported by NASA under grant or cooperative agreement award number NNX17AJ31G. This work was also supported by a NASA Space Technology Research Fellowship (NASA grant number 80NSSC18K1157). This work is supported by the Translational Research Institute through NASA Cooperative Agreement NNX16AO69A. Any opinions, findings, and conclusions or recommendations expressed in this material are those of the author(s) and do not necessarily reflect the views of the National Aeronautics and Space Administration (NASA) or the Translational Research Institute for Space Health (TRISH).

Illustrative figures in part created with BioRender.com.

The authors report no conflicts of interest.

## Abbreviations

CHM: pre-packed chromatography
EHS: environmental, health, and safety
ELP: elastin-like polypeptide
ESM: equivalent system mass
Fc: fragment crystallizable
ISRU: *in situ* resource utilization
ISS: International Space Station
mAb: monoclonal antibody
MAG: magnetic bead
PMI: process mass intensity
OLE: oilbody-oleosin
RMA: reference mission architecture
SPN: spin column
VIN: plant virus-based nanoparticle
xESM: extended equivalent system mass.

