## Supplementary Information for "Evaluating the cost of pharmaceutical purification for a long-duration space exploration medical foundry"

***Supplementary Material***

**Supplementary Table S1.** Example commercially approved monoclonal antibody (mAb) therapies of relevance to human health in space that have been considered in the determination of the reference mission pharmaceutical demand. Asterisk (*) denotes an antibody drug conjugate.

| mAb | Indication | Dose | Need Basis |
|---|---|---|---|
| Erenumab-aooe (FDA Label) | Migraine headache prevention | 70 mg (or 140 mg) | 1 dose/month |
| Romosozumab (FDA Label) | Bone regeneration | 210 mg | 1 dose/month |
| Gemtuzumab ozogamicin* (FDA Label) | Acute myeloid leukemia | 6 $mg/m^2$; 3 $mg/m^2$; 2 $mg/m^2$ | day 1/day 8/every 4 weeks; 1 course/year |

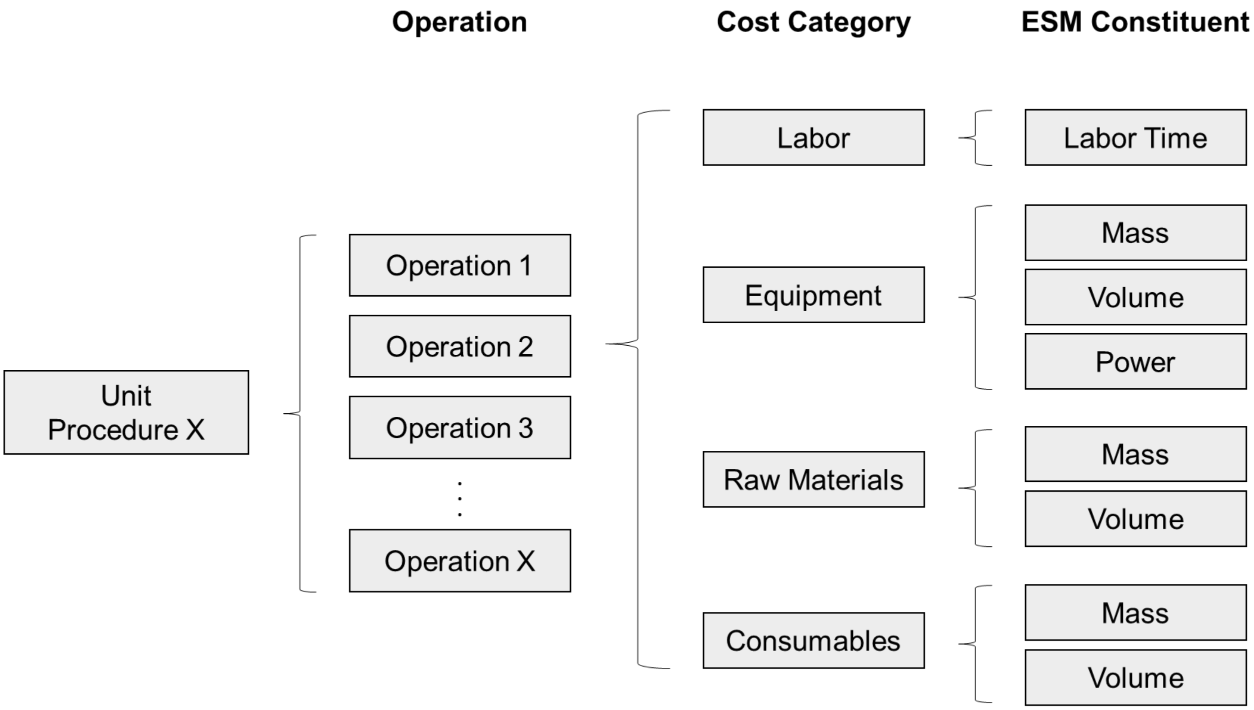


**Supplementary Figure S1.** Schematic of deterministic unit procedure model construction grouped by operation, cost category, and equivalent system mass (ESM) constituent.

**Supplementary Table S2.** Unit procedure assumptions for maximal feed stream volume and monoclonal antibody (mAb) binding capacity.

| Unit Procedure | Code | mAb Binding Capacity | Maximal Feed Stream Volume |
|---|---|---|---|
| Pre-packed chromatography | CHM | 30 mg/mL resin | N/A |
| Spin column | SPN | 1 mg/column | 0.6 mL |
| Magnetic bead | MAG | 27 mg/mL bead slurry | 0.3 mL |
| Plant virus-based nanoparticle | VIN | 4 mg/mL stock solution | 2 mL* |
| Elastin-like polypeptide | ELP | 0.42 mg/mL stock solution | 0.8 mL*,[¥] |
| Oilbody-oleosin | OLE | 6.74 mg/mL stock solution | 0.1 mL*,[γ] |

* based on 2 mL unit volume; actual feed stream volume added is based on the amount of stock solution required and thus mAb quantity in the feed stream.

[¥] reduced from 2 mL maximal to account for volume needed for salt solution addition (0.4 mL) and required 1:1 volume ratio of ELP:mAb.

γ reduced from 2 mL maximal to account for required 1:20 volume ratio of OLE:mAb.

**Supplementary Table S3.** Labor time standardizations applied to common operations across unit procedures.

| Operation | Value | Unit |
|---|---|---|
| Monitoring | 0.05 | labor hour/hour |
| Preparation (incubation + centrifugation) | 1.0 | min/effective batch |
| Pipetting liquid | 0.5 | min/solution type |
| | 0.1 | min/additional unit/effective batch |
| Resuspending pellet | 1 | min/unit |

**Supplementary Table S4.** Equivalency factor values used to generate extended equivalent system mass values from constituents of mass, volume, power, cooling, and labor.

| Segment | $L_{eq}$ (kg/kg) | $M_{eq}$ (kg/kg) | $V_{eq}$ (kg/m$^3$) | $P_{eq}$ (kg/kW) | $C_{eq}$ (kg/kW) | $T_{eq}$ (kg/CM-h) |
|---|---|---|---|---|---|---|
| Pre-deployment (Pd) | 2.77 | 1 | 9.16 | 237 | 40 | 0.7 |
| Transit to Mars (Tr1) | 10 | 1 | 133.8 | 136 | 50 | 0.7 |
| Surface Operation (Su) | 1 | 1 | 9.16 | 228 | 145 | 0.7 |
| Return Transit (Tr2) | 10 | 1 | 133.8 | 136 | 50 | 0.7 |

**Supplementary Table S5.** Optimal number of effective batches per mission in the base case scenario for each unit procedure, as determined via minimization of equivalent system mass.

| Unit procedure | Optimal number of batches per mission |
|---|---|
| CHM | 342 |
| SPN | 948 |
| MAG | 5670 |
| VIN | 360 |
| ELP | 2058 |
| OLE | 846 |

**Supplementary Table S6.** Mars surface mission equivalency factor values used by Zabel, 2020 of a space greenhouse.

| Segment | $M_{eq}$ (kg/kg) | $V_{eq}$ (kg/m$^3$) | $P_{eq}$ (kg/kW) | $C_{eq}$ (kg/kW) | $T_{eq}$ (kg/CM-h) |
|---|---|---|---|---|---|
| Surface Operation (Su) | 1.0 | 215.5 | 87.0 | 146.0 | 0.465 |

**Supplementary Table S7.** Optimal number of effective batches per mission in the mAb stream composition scenario analysis conditions for each unit procedure, as determined via minimization of equivalent system mass.

| mg/mL | 0.10 | 0.20 | 0.35 | 0.50 | 075 | 1.00 | 1.50 | 2.00 | 5.00 |
|---|---|---|---|---|---|---|---|---|---|
| CHM | 342 | 342 | 342 | 342 | 342 | 342 | 342 | 342 | 342 |
| SPN | 9450 | 4728 | 2700 | 1896 | 1260 | 948 | 630 | 568 | 568 |
| MAG | 56694 | 28350 | 16200 | 11340 | 7560 | 5670 | 3780 | 2838 | 1134 |
| VIN | 2910 | 1494 | 882 | 639 | 450 | 360 | 264 | 216 | 132 |
| ELP | 7092 | 3546 | 2058 | 2058 | 2058 | 2058 | 2058 | 2058 | 2058 |
| OLE | 2988 | 1494 | 854 | 846 | 846 | 846 | 846 | 846 | 846 |

**Supplementary Table S8.** Optimal number of batches per mission in the mAb demand scenario analysis conditions for each unit procedure, as determined via minimization of equivalent system mass.

| Demand x $10^3$ (mg mAb/mission) | 1.0 | 2.0 | 3.5 | 5.0 | 7.5 | 10.0 | 15.0 | 20.0 | 30.0 |
|---|---|---|---|---|---|---|---|---|---|
| CHM | 36 | 70 | 120 | 172 | 257 | 342 | 512 | 682 | 1022 |
| SPN | 96 | 192 | 336 | 474 | 714 | 948 | 1422 | 1890 | 2844 |
| MAG | 568 | 1134 | 1988 | 2838 | 4260 | 5670 | 8508 | 11340 | 17010 |
| VIN | 36 | 72 | 126 | 180 | 270 | 360 | 533 | 714 | 1068 |
| ELP | 210 | 414 | 726 | 1032 | 1548 | 2058 | 3090 | 4116 | 6174 |
| OLE | 86 | 170 | 300 | 422 | 632 | 846 | 1264 | 1692 | 2532 |

**Supplementary Table S9.** List of centrifuge models used in the alternative centrifuge scenario.

| Model | Vendor | Capacity | Mass (kg) | Dimensions (cm) | Power (kW) |
|---|---|---|---|---|---|
| MiniSpin | Eppendorf | 12 | 3.7 | 22.5 x 23.0 x 13.0 | 0.085 |
| 5418R | Eppendorf | 18 | 22 | 0.0345 | 0.320 |
| 5427R | Eppendorf | 48 | 30 | 31.9 x 54.0 x 25.4 | 0.550 |

**Supplementary Table S10.** Optimal number of effective batches per mission in the centrifuge model alternative scenario conditions for each analyzed unit procedure, as determined via minimization of equivalent system mass.

| Centrifuge model | MiniSpin | 5418R | 5427R |
|---|---|---|---|
| SPN | 1422 | 948 | 360 |
| VIN | 533 | 360 | 137 |
| ELP | 3090 | 2058 | 774 |
| OLE | 1264 | 846 | 317 |
